# Successful retrieval is its own reward

**DOI:** 10.1101/2024.07.26.605274

**Authors:** Devyn E. Smith, Nicole M. Long

## Abstract

The ability to successfully remember past events is critical to our daily lives, yet the neural mechanisms underlying the motivation to retrieve is unclear. Although reward-system activity and feedback-related signals have been observed during memory retrieval, whether this signal reflects intrinsic reward or goal-attainment is unknown. Adjudicating between these two alternatives is crucial for understanding how individuals are motivated to engage in retrieval and how retrieval supports later remembering. To test these two accounts, we conducted between-subjects recognition memory tasks on human participants undergoing scalp electroencephalography and varied test-phase goals (recognize old vs. detect new items). We used an independently validated feedback classifier to measure positive feedback evidence. We find positive feedback following successful retrieval regardless of task goals. Together, these results suggest that successful retrieval is intrinsically rewarding. Such a feedback signal may promote future retrieval attempts as well as bolster later memory for the successfully retrieved events.

## Introduction

The ability to accurately remember people whom you have met and places that you have visited is highly advantageous, enabling basic survival, supporting social interactions, and giving rise to self-identity. Yet the mechanisms that motivate individuals to engage in retrieval are not well understood. Although correctly remembering can lead to downstream rewards–e.g. remembering which patch of woods has the most berries will promote subsequent food-related rewards–it might be beneficial for the act of retrieving itself to produce reward signals. Prior work has shown that remembering positive autobiographical memories is accompanied by positive feelings and reward system activation^1^, which suggests that successful retrieval may be intrinsically rewarding. However, as accomplishing task goals leads to rewards, to the extent that successful retrieval satisfies a task goal, retrieval-related reward signals may instead reflect goal-attainment. It is important to determine the driver of retrieval-related reward as the tendency to retrieve–and possibly the strategies used to encode and retrieve memories–may be impacted by reward signals during successful retrieval. The aim of this study was to investigate whether test-phase neural signals reflect intrinsic reward or goal-attainment.

Converging evidence from non-invasive human functional magnetic resonance imaging (fMRI) and scalp electroencephalography (EEG) recordings has shown reward-system and feedback-related signals during memory retrieval^2^. Striatal activity is greater during successful item retrieval (hit) compared to correct identification of a new stimulus (correct rejection, CR;^3–6^). The striatum has long been linked with decision-making, representation of reward, and motivation in the context of extrinsic rewards–external rewards delivered to the individual such as money, points, or positive feedback^7, 8^. Given that extrinsic rewards are rarely provided during recognition memory tests, there is no *a priori* reason to anticipate striatal engagement during retrieval and suggests the existence of a post-retrieval reward signal, yet due to the low temporal resolution of fMRI, when these signals occur relative to retrieval is unclear. We have recently shown that frontocentral theta power (4-8 Hz) decreases following successful retrieval^9^. Frontocentral theta is engaged during decision-making and cognitive control tasks and decreases following positive feedback and correct responses^10–12^. Theta power increases have been linked with the error- and feedback-related negativity event-related potential components following incorrect responses^13–17^. The relative decrease in theta power that we find following hits compared to CRs may be analogous to the relative decrease in theta power following positive relative to negative outcomes. Theta power may reflect a non-specific mechanism that coordinates “internal and external performance relevant information.”^12^. Critically, whereas fMRI cannot elucidate precisely when feedback responses occur, our scalp EEG findings clearly demonstrate that this theta power decrease occurs *after* the memory response is made. A post-response, hit-specific theta power decrease is consistent with the interpretation that a decision-making-related intrinsic feedback signal follows successful retrieval, yet a direct test of this account is lacking.

Despite abundant evidence for feedback-related signals during memory tasks, such signals may reflect goal-attainment rather than a response specific to successful retrieval. In a typical recognition memory task, the participants’ goal is to recognize old items, thus successful retrieval is confounded with task goals. Evidence for the goal-attainment account stems from recognition studies with extrinsic rewards^18^. When participants have the potential to earn an extrinsic reward for CRs, striatal activity is greater in response to CRs than hits^18^. However, extrinsic rewards are known to influence intrinsic rewards^19^. Therefore it is unclear whether feedback-related signals during successful retrieval are similarly modulated by goals in the absence of extrinsic rewards.

Our hypothesis is that successful retrieval is intrinsically rewarding. The alternative is that feedback-related signals following successful retrieval reflect goal-attainment. To adjudicate between these hypotheses, we conducted two pre-registered recognition memory scalp EEG experiments (E1, E2) in which we manipulated test-phase goals. Participants’ goal was to either recognize old items (E1) or detect new items (E2). We measured post-response theta power in two frontocentral regions of interest. To directly assess post-response feedback processing, we trained whole-brain multivariate pattern classifiers to distinguish correct from incorrect trials in two flanker tasks. We applied this independent decoder to the recognition data to avoid informal reverse inference^20^ and measure test-phase feedback. All analyses were pre-registered except where indicated. To the extent that successful retrieval is intrinsically rewarding, we should find the same positive post-response feedback signals following hits regardless of memory goals.

## Results

### Post-response theta power decreases for hits vs CRs regardless of goals

We have previously demonstrated with these data^21^ that the task goal manipulation (E1: recognize old vs. E2: detect new) impacts recognition memory responses whereby reaction times are faster when a participant’s goal (e.g. detect new) matches the probe type (e.g. lure; see Figure 2B in the referenced manuscript). This finding is in line with the encoding specificity hypothesis^22^ whereby performance is better for matched, compared to mismatched, study and test-phase contexts and indicates that the goal manipulation was sufficient to modulate behavior.

Our first objective was to independently replicate our prior finding^9^ that theta power is modulated by successful retrieval leading up to and following memory responses (Figure 1). To specifically test for a pre- vs. post-response dissociation in theta power for hits and CRs across E1 and E2, we averaged signals within the 500 ms pre- and post-response time intervals (Figure 1, insets). To the extent that theta power is modulated by successful retrieval, the same post-response signals should dissociate hits vs. CRs regardless of memory goals. However, to the extent that theta power dissociations are driven by goal-attainment, we would expect decreased post-response theta for hits compared to CRs in E1 and decreased post-response theta for CRs compared to hits in E2. We conducted two 2*×*2*×*2 mixed effects ANOVAs (meANOVAs, Table 1), one for each of two frontocentral ROIs (left central; LC, right central; RC) with factors of experiment (E1, E2), time interval (pre-response, post-response) and response type (hit, CR). If successful retrieval drives the post-response theta decrease for hits, we would expect to find an interaction between response type and time interval and no experiment driven interactions. Alternatively, if goal-attainment drives the post-response theta decrease, we would expect to find an interaction between experiment, time interval, and response type. For both ROIs, we find a significant interaction between response type and time interval. In LC, this interaction was driven by greater theta power for hits (M=0.28, SD=0.29) relative to CRs (M=0.19, SD=0.25) in the pre-response time interval (*t* _71_=2.896, *p*=0.005, *d* =0.3254) and greater theta power for CRs (M=0.13, SD=0.25) relative to hits (M=- 0.01, SD=0.29) in the post-response time interval (*t* _71_=4.984, *p<*0.0001, *d* =0.5216). In RC, this interaction was driven by no difference in theta power for hits (M=0.23, SD=0.28) relative to CRs (M=0.21, SD=0.25) in the pre-response time interval *t* _71_=0.6338, *p*=0.5283, *d* =0.0698) and greater theta power for CRs (M=0.19, SD=0.24) relative to hits (M=0.005, SD=0.28) in the post-response time interval (*t* _71_=6.216, *p<*0.0001, *d* =0.7032). The three-way interaction between response type, time interval, and experiment was not significant. Bayes factor analysis revealed that a model without the three-way interaction term (H_0_) is preferred to a model with the three-way interaction term (H_1_) in both ROIs (LC, BF_10_=0.27, moderate evidence for H_0_; RC, BF_10_=0.27, moderate evidence for H_0_). Together, these results indicate that theta power is modulated by successful retrieval following memory responses regardless of task goals and is specifically consistent with the interpretation that successful retrieval produces a positive feedback signal represented by relative theta power decreases.

**Figure 1.**
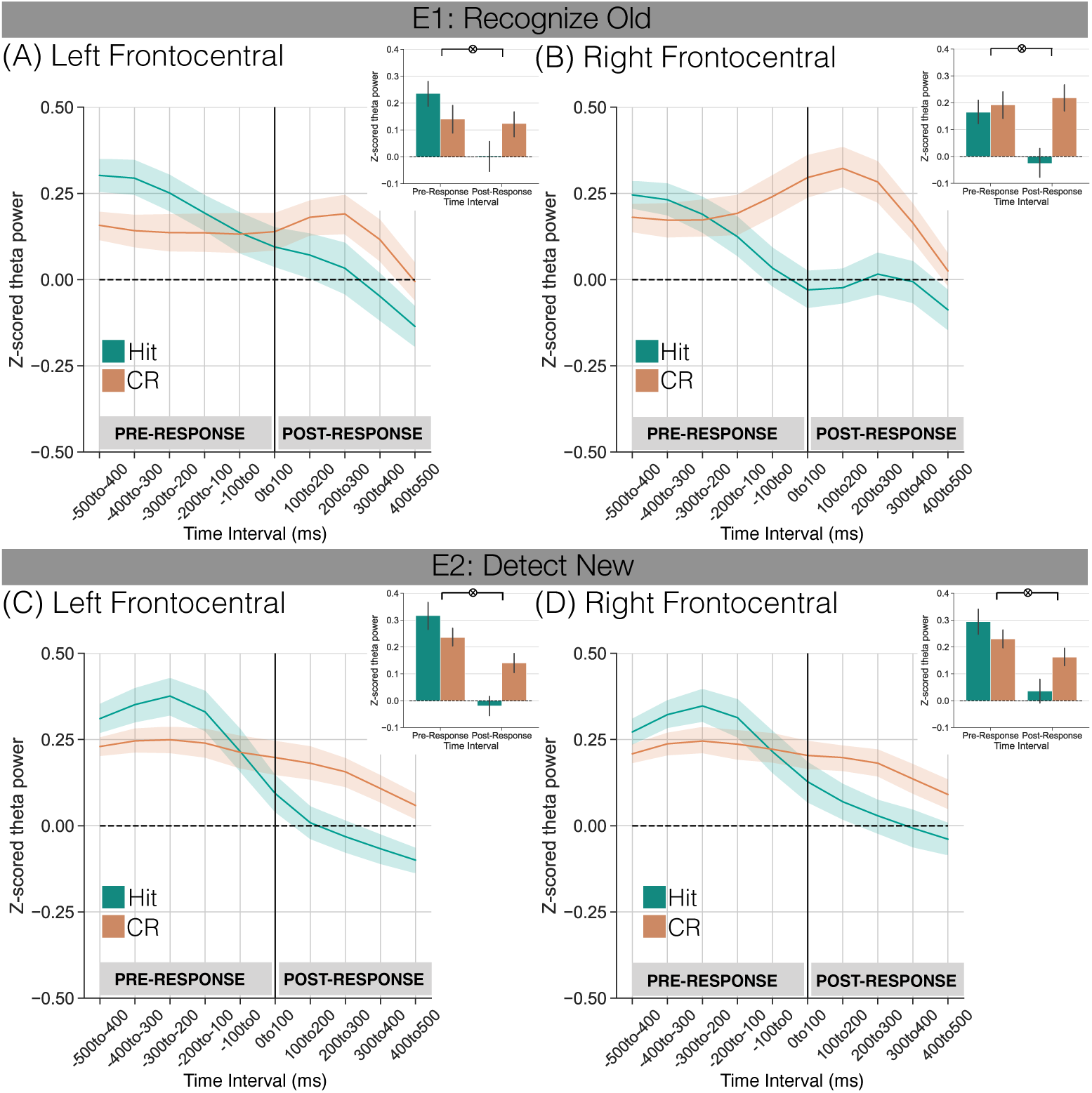
Theta power dissociations across hits and correct rejections preceding and following memory responses in E1 and E2. Response-locked z-transformed theta power (4-8 Hz) for the left and right central ROIs. The solid vertical black line indicates when the response was made. Hits are shown in teal, correct rejections (CRs) are shown in orange. Error bars reflect standard error of the mean. **(A-B)** E1, participants recognize old items. **(A)** Over left central ROI, we find a significant interaction between response type and time interval (p *<* 0.001) driven by numerically greater pre-response theta power for hits than CRs and greater post-response theta power for CRs than hits. **(B)** Over right central ROI, we find a significant interaction between response type and time interval (p *<* 0.001) driven by greater post-response theta power for CRs than hits. **(C-D)** E2, participants detect new items. **(C)** Over left central ROI, we find a significant interaction between response type and time interval (p *<* 0.001) driven by greater pre-response theta power for hits than CRs and greater post-response theta power for CRs than hits. **(D)** Over right central ROI, we find a significant interaction between response type and time interval (p *<* 0.001) driven by numerically greater pre-response theta power for hits than CRs and greater post-response theta power for CRs than hits.

**Table 1.**
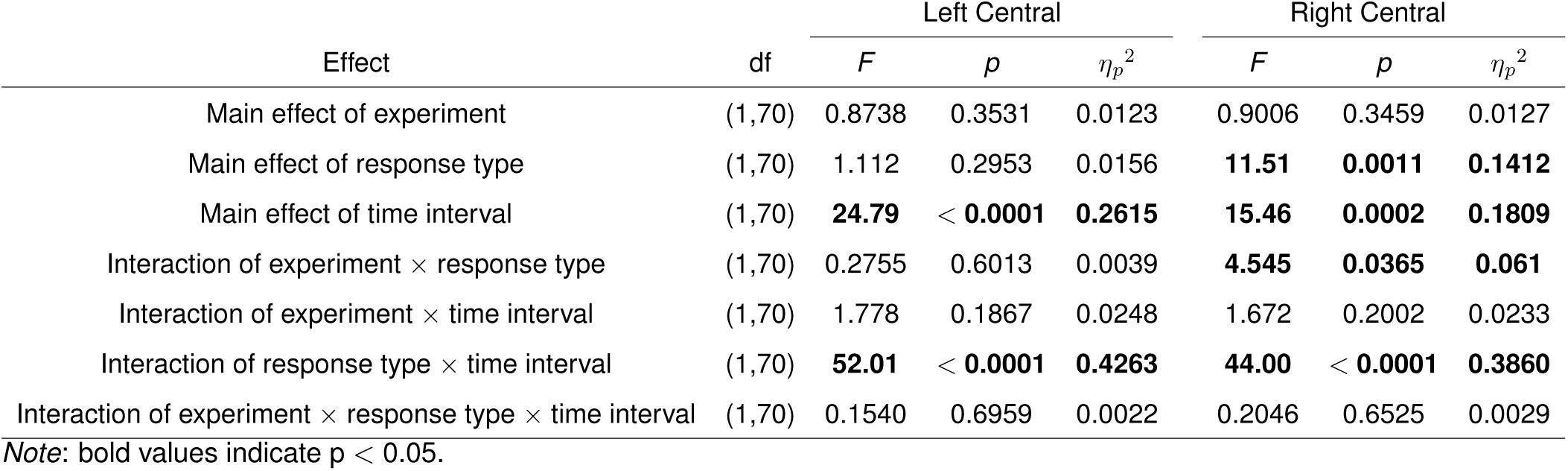
Mixed effects ANOVAs assessing theta power for left and right central ROIs as a function of experiment, response type, and time interval.

|  |  | Left Central |  |  | Right Central |  |  |
|---|---|---|---|---|---|---|---|
| Effect | df | *F* | *p* | *η<sub>p</sub>*<sup>2</sup> | *F* | *p* | *η<sub>p</sub>*<sup>2</sup> |
| Main effect of experiment | (1,70) | 0.8738 | 0.3531 | 0.0123 | 0.9006 | 0.3459 | 0.0127 |
| Main effect of response type | (1,70) | 1.112 | 0.2953 | 0.0156 | **11.51** | **0.0011** | **0.1412** |
| Main effect of time interval | (1,70) | **24.79** | < **0.0001** | **0.2615** | **15.46** | **0.0002** | **0.1809** |
| Interaction of experiment × response type | (1,70) | 0.2755 | 0.6013 | 0.0039 | **4.545** | **0.0365** | **0.061** |
| Interaction of experiment × time interval | (1,70) | 1.778 | 0.1867 | 0.0248 | 1.672 | 0.2002 | 0.0233 |
| Interaction of response type × time interval | (1,70) | **52.01** | < **0.0001** | **0.4263** | **44.00** | < **0.0001** | **0.3860** |
| Interaction of experiment × response type × time interval | (1,70) | 0.1540 | 0.6959 | 0.0022 | 0.2046 | 0.6525 | 0.0029 |

*Note*: bold values indicate p < 0.05.

### Robust feedback decoding in the flanker task

Before testing our central hypothesis, we validated that feedback and response accuracy in the flanker task could be reliably decoded. We conducted multivariate pattern classification analyses in which we trained a classifier to discriminate positive versus negative feedback or correct versus incorrect trials based on a feature space composed of all 63 electrodes and 46 frequencies ranging from 2-100 Hz. For this analysis, we averaged z-power over the 100 ms preceding the response. Using leave-one-participant-out cross-validated classification (penalty parameter=0.0001), mean classification accuracy was significantly greater than chance for both the feedback (M=59.42%, SD=9.520%; *t* _68_=8.171, *p<*0.0001, *d* =1.401; Figure 2A, top row) and no-feedback (M=64.85%, SD=10.16%; *t* _14_=5.483, *p*=0.0001, *d* =2.083; Figure 2A, bottom row) experiment.

**Figure 2.**
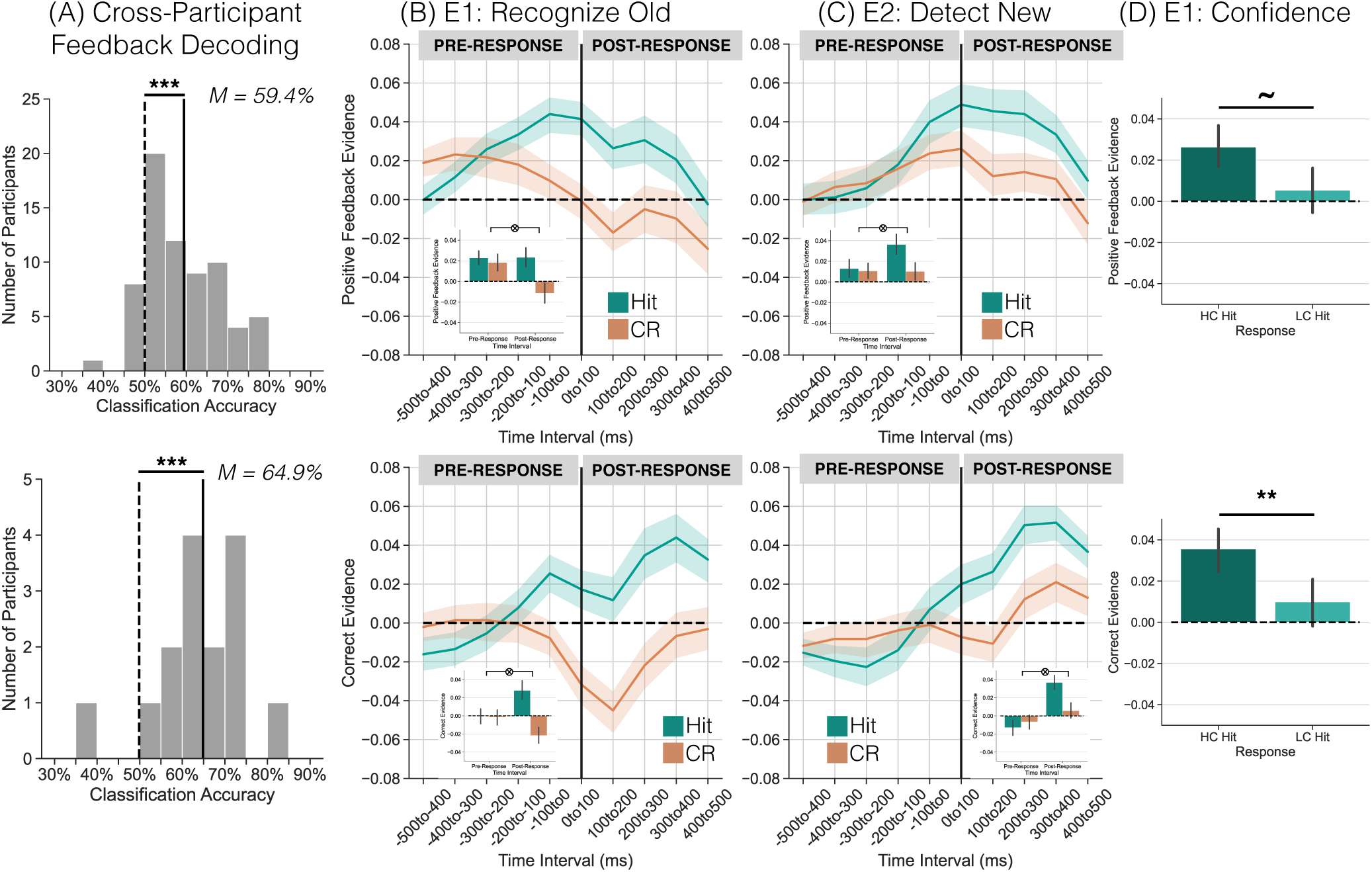
Positive feedback and correct evidence preceding and following memory responses in E1 and E2. The top row shows results from the feedback trained classifier; the bottom row shows results from the no- eedback trained classifier. **(A)** Mean cross-participant classification accuracy across all participants (solid vertical line) is shown along with a histogram of classification accuracies for individual participants (gray bars) and mean classification accuracy for permuted data across all participants (dashed vertical line). Mean positive feedback classification accuracy was 59.4% (top row) and mean correct feedback classification accuracy was 69.4% (bottom row), both of which differed significantly from chance (two-tailed, paired t-test, *p* =*<* 0.0001). **(B-C)** Positive y-axis values indicate greater positive feedback/correct evidence. The solid vertical line at time 0 to 100 ms indicates the response. Each panel shows positive feedback/correct evidence separated by response type (teal: hit; orange: CR) across the 500 ms pre-response and post-response time intervals. There is greater post-response positive feedback/correct evidence for hits compared to CRs (*p <* 0.0001), regardless of experiment goals. **(D)** Positive feedback/correct evidence is modulated by confidence following memory responses in E1. We assessed post-response (0 to 500 ms) positive feedback/correct evidence for E1 high confident (HC; dark teal) and low confident (LC; light teal) responses. There is greater positive feedback evidence (*p* = 0.0510) and correct evidence (*p* = 0.0068) for HC hits compared to LC hits. Error bars represent standard error of the mean. *∼ p <* 0.10, \*\**p <* 0.01, \*\*\**p <* 0.001.

### Greater positive feedback evidence for hits vs CRs regardless of goals

Our central goal was to test the hypothesis that successful retrieval is intrinsically rewarding. Our interpretation is that post-response theta power dissociations reflect a feedback signal whereby positive feedback is greater following hits compared to CRs. However, participants did not receive any explicit feedback during E1 and E2, therefore we cannot definitively conclude that these post-response signals reflect positive feedback. In order to test our hypothesis, we developed an independent measure of feedback by training a classifier on pre-response feedback signals in a flanker task. We then tested the classifier on test-phase hits and CRs in E1 and E2. To the extent that successful retrieval is intrinsically rewarding, we expect to find greater positive feedback evidence following hits compared to CRs in both experiments. Alternatively, to the extent that feedback signals reflect goal-attainment, we expect to find greater positive feedback evidence following CRs compared to hits in E2 relative to E1.

We measured feedback evidence across ten 100 ms time intervals from 500 ms preceding to 500 ms following the test-phase response in E1 and E2 (Figure 2B,C, top row). To specifically test for a pre- vs. post-response dissociation in positive feedback evidence following hits and CRs across E1 and E2, we averaged signals within the 500 ms pre- and post-response time intervals (Figure 2B,C insets, top row). We conducted a 2*×*2*×*2 meANOVA (Table 2) with factors of experiment (E1, E2), time interval (pre-response, post-response) and response type (hit, CR). We find a significant interaction between time interval and response type driven by no difference in positive feedback evidence for hits (M=0.0180, SD=0.0472) relative to CRs (M=0.0145, SD=0.0487) in the pre-response time interval (*t* _71_=0.7394, *p*=0.4621, *d* =0.0719) and significantly greater positive feedback evidence for hits (M=0.0298, SD=0.0561) relative to CRs (M=-0.0007, SD=0.0552) in the post-response time interval (*t* _71_=5.952, *p<*0.0001, *d* =0.5485). The three-way interaction between response type, time interval, and experiment was not significant. Bayes factor analysis revealed that a model without the three-way interaction term (H_0_) is preferred to a model with the three-way interaction term (H_1_; BF_10_=0.27, moderate evidence for H_0_). These results support the interpretation that successful retrieval is intrinsically rewarding.

**Table 2.** Mixed effects ANOVAs assessing positive feedback and correct evidence as a function of experiment, response type, and time interval.

|  |  | Positive Feedback Evidence |  |  | Correct Evidence |  |  |
|---|---|---|---|---|---|---|---|
| Effect | df | *F* | *p* | *η<sub>p</sub>*<sup>2</sup> | *F* | *p* | *η<sub>p</sub>*<sup>2</sup> |
| Main effect of experiment | (1,70) | 0.1966 | 0.6588 | 0.0028 | 0.2227 | 0.6385 | 0.0032 |
| Main effect of response type | (1,70) | **17.01** | **0.0001** | **0.1955** | **22.90** | < **0.0001** | **0.2465** |
| Main effect of time interval | (1,70) | 0.0828 | 0.7743 | 0.0012 | **9.621** | **0.0028** | **0.1208** |
| Interaction of experiment × response type | (1,70) | 0.4502 | 0.5044 | 0.0064 | 2.670 | 0.1067 | 0.0367 |
| Interaction of experiment × time interval | (1,70) | **5.053** | **0.0277** | **0.0673** | **5.639** | **0.0203** | **0.0746** |
| Interaction of response type × time interval | (1,70) | **25.22** | < **0.0001** | **0.2649** | **62.73** | < **0.0001** | **0.4726** |
| Interaction of experiment × response type × time interval | (1,70) | 0.3481 | 0.5571 | 0.0049 | 1.003 | 0.3199 | 0.0141 |

*Note*: bold values indicate p < 0.05.

To validate that the hit-specific feedback signal we observe is not a product of the visually presented feedback stimuli and/or external, extrinsic rewards, we collected data in a second flanker task with no-feedback. We used the same approach as in the feedback condition, except that flanker trials were coded on the basis of accuracy rather than feedback (Figure 2B,C insets, bottom row). We conducted an exploratory 2*×*2*×*2 meANOVA (Table 2) with factors of experiment (E1, E2), time interval (pre-response, post-response) and response type (hit, CR). We find a significant interaction between time interval and response type driven by no difference in correct evidence for hits (M=-0.0067, SD=0.0504) relative to CRs (M=-0.0041, SD=0.0479) in the pre-response time interval (*t* _71_=0.5512, *p*=0.5833, *d* =0.0527) and significantly greater correct evidence for hits (M=0.0325, SD=0.0559) relative to CRs (M=-0.0080, SD=0.0539) in the post-response time interval (*t* _71_=8.128, *p<*0.0001, *d* =0.7376). The three-way interaction between response type, time interval, and experiment was not significant. Bayes factor analysis revealed that a model without the three-way interaction term (H_0_) is preferred to a model with the three-way interaction term (H_1_; BF_10_=0.29, moderate evidence for H_0_).

We confirmed that these hit-related effects do not reflect either objective or subjective “oldness” by comparing positive feedback and correct evidence following error trials (misses, false alarms) to hits. Hits show significantly more positive feedback/correct evidence than all other response types (Supplemental Material). We further verified that these findings are unlikely to be the result of response conflict or cognitive control by showing that although there is a congruency dissociation between hits and CRs, it is driven by greater pre-response congruency evidence for CRs relative to hits across both experiments (Supplemental Material). These results support the interpretation that successful retrieval is intrinsically rewarding.

### Positive feedback evidence follows high confidence hits

Greater positive feedback signals following hits compared to CRs implies that participants are aware that they have successfully retrieved, yet participants received no explicit feedback regarding their performance. We can gain some insights by investigating the extent to which feedback/correct evidence is modulated by response confidence on hits in E1. By definition, participants are more confident that they are correct when they give a high confidence response. Therefore, positive feedback/correct evidence should be greater following high, compared to low, confidence hits. We conducted an exploratory analysis in which we assessed post-response (0 to 500 ms) positive feedback evidence following high and low confidence hits (Figure 2D, top row). We find marginally greater positive feedback evidence for high (M=0.0263, SD=0.0594) compared to low (M=0.0054, SD=0.0678) confidence hits (*t* _35_=2.021, *p*=0.0510, *d* =0.3291). We repeated this analysis for the no-feedback flanker training data with correct evidence as the dependent variable (Figure 2D, bottom row). We find significantly greater correct evidence for high (M=0.0356, SD=0.0646) compared to low (M=0.0099, SD=0.0695) confidence hits (*t* _35_=2.879, *p*=0.0068, *d* =0.3828). Thus, when participants have high confidence that they are correct, and they have correctly remembered a past experience, we find greater positive feedback/correct evidence than when participants are less confident in their response.

## Discussion

The aim of this study was to identify whether test-phase feedback-related signals reflect successful retrieval or goal-attainment. We conducted two independent recognition memory experiments in which we manipulated participants’ test-phase goals to either recognize old items (E1) or detect new items (E2) and did not provide any explicit feedback. We recorded scalp EEG and used a cross-study decoding approach^21, 23^ to measure response-locked positive feedback evidence during the test-phase of each experiment. Across both experiments, we find post-response theta power decreases and greater positive feedback/correct evidence for hits compared to CRs. Together, these findings suggest that successful retrieval is intrinsically rewarding. The implication is that such feedback signals may motivate both future retrieval attempts and the strategies used to encode and retrieve memories.

Following hits–successful retrieval–we find a decrease in frontocentral theta power and an increase in positive feedback/correct evidence relative to CRs. Substantial prior work has shown that striatal activity is greater for hits relative to CRs during recognition memory tests^3–6, 24–26^. The striatum is well known to respond to extrinsic rewards such as money^27, 28^. That the striatum shows increased activation selectively for hits suggests that successful retrieval, correctly remembering a past experience, may be intrinsically rewarding. Such an account is consistent with the finding that remembering positively-valenced past experiences is associated with positive feelings and reward system activity^1^. However, it was an open question as to when these signals occur relative to the memory response. By leveraging the high temporal resolution of scalp EEG, we specifically find post-response feedback/accuracy-related dissociations between hits and CRs. Our interpretation is that following memory-related processes such as search and evidence accumulation^29, 30^, individuals conduct a post-response process in which they evaluate the just-made memory response. Our results suggest a positive feedback signal following successful retrieval.

We find evidence for positive feedback/correct signals following successful retrieval regardless of task goals. In a typical recognition memory experiment, participants’ goal is to recognize old items and no extrinsic feedback is provided. Thus, post-response theta power decreases and positive evidence increases may reflect a response to goal-attainment, rather than to successful retrieval^31^. The striatum shows elevated activation in response to CRs relative to hits when CRs are linked with extrinsic rewards^18^, suggesting that feedback-related signals may be a response to achieving the goals set by the experimenter (recognize old or detect new), rather than successful retrieval per se. However, although extrinsic rewards enable experimenter control, their introduction is likely to impact intrinsic processing as extrinsic rewards can undermine or modify intrinsic motivation and alter satisfaction with tasks^19^. Explicit feedback can modulate how information is processed^32^ and also has variable utility depending on the information being learned^33^. Thus, to test the goal-attainment account while not introducing extrinsic rewards, we combined a subtle experimental manipulation of task goals with a novel approach in which we developed a feedback-related multivariate pattern classifier. We instructed participants to either recognize old items (E1) or detect new items (E2), a manipulation that was sufficient to induce reaction time changes to test-phase responses^21^. Applying the feedback classifier to the test-phase neural data enabled us to assess evidence of positive feedback following hits and CRs. Although both hits and CRs constitute correct responses–meaning that both response types could yield a positive feedback signal–and despite the lack of extrinsic feedback, we find positive feedback selectively following hits. Furthermore, we find this pattern even when participants’ goal is to detect new items–that is, despite CRs being relevant to the task goals in E2, they still show less positive feedback evidence relative to hits. We replicate these findings when we utilized a training set derived from a no-feedback flanker task. We find greater correct evidence following hits relative to CRs across both experiments. Together, these findings provide strong evidence for the account that successful retrieval is intrinsically rewarding.

Our investigation of post-response feedback signals began with our prior observation of greater post-response theta power for CRs relative to hits^9^; however, other neural signals have been linked with feedback, error monitoring, and reward processing^12, 17^. Specifically, decision-making tasks have revealed error- and feedback-related negativities (ERN/FRN,^34, 35^) as well as the reward positivity, a central positive event-related potential (ERP) component that emerges around 200-300 ms following feedback^36, 37^. We chose not to utilize ERPs as the training signal for this study given that ERPs depend on precise time-locking to the onset of a stimulus or response. Although we would anticipate timing consistency across traditional decision-making tasks, we did not expect that the time course of processing should unfold along the same time frame in the current recognition studies. Thus spectral activity, which captures both time-locked and non-time-locked signals, are better suited to the present investigation. Given the close link between the ERN, FRN and theta^16, 38^, we expect that these components are likely captured in our spectral classifiers. Critically, our feedback/accuracy classifiers incorporate features beyond central theta power, the original starting point of the work. The features cover spectral frequencies from 2-100Hz and include all electrodes. The rationale for this inclusion is both practical–cross participant classification with whole brain/all frequencies outperformed classification restricted to central theta–and conceptual, in that whole brain/all frequency classifiers will capture multiple aspects of reward, error, and feedback processing. Thus, we are confident that these classifiers are detecting such signals following successful retrieval.

How do participants ‘know’ that they have successfully retrieved? Since no feedback is provided in the current experiments–or in most recognition experiments–participants do not receive any external confirmation that they are correct. Yet, anecdotally, it is possible to ‘feel’ that one has correctly retrieved. To manipulate test-phase goals without introducing substantial other experimental differences, we kept test-phase reporting to a minimum. However, the confidence ratings in E1 provide some insights into the mapping between successful retrieval and positive feedback. High confidence responses may reflect reinstatement of source or contextual details^39, 40^. We find greater positive feedback/correct evidence following high, compared to low, confidence hits, consistent with the interpretation that when participants have more and/or stronger information upon which to make a decision, a greater positive feedback signal is generated. In paradigms with explicit feedback, more information produces greater caudate responses^41^. It may thus be that positive feedback signals are only emitted when participants are highly confident in their retrieval; however, future studies will be needed to fully characterize the scope of positive feedback following memory responses.

Intrinsic signals following successful retrieval have the potential to substantially impact memory and behavior. First, such signals may promote future retrieval attempts. Likewise, deficits in reward and feedback signaling found in clinical conditions^42–44^ may explain co-morbid memory deficits whereby a lack of positive feedback in response to successful retrieval may diminish future retrieval attempts. Second, to the extent that test-phase signals mirror the effects found for motivated encoding^2, 45–49^, increases in test-phase positive feedback may enhance subsequent memory for successfully retrieved items. Extrinsic post-response rewards facilitate subsequent memory^50^. Intrinsic signaling following successful retrieval may enable the reinforcement of both the memories themselves and the strategies used to encode and retrieve those memories. For instance, prior work has shown that the task context in which an item was encoded is reinstated during test^51^. Insofar as test-phase feedback signals reinforce the contents of retrieval, that task context will also be reinforced. The consequences of such a reinforcement mechanism is that if participants use and retrieve suboptimal memory strategies (such as a shallow orienting task), maladaptive strategies may be reinforced. Similarly, participants’ belief about their response correctness–related to the points in the paragraph above–may interact with the reinforcement of correct vs. incorrect information. Future work will be needed to test these accounts.

In conclusion, we find evidence for positive/correct feedback following successful retrieval regardless of task goals. These results support the interpretation that successful retrieval is intrinsically rewarding. The consequences of intrinsically rewarding successful retrieval may extend far beyond the current moment to influence both future remembering of the retrieved experience and how future experiences may be remembered.

## Supporting information

Supplemental Results

## Resource availability

### Lead Contact

Requests for further information and resources should be directed to and will be fulfilled by the lead contact, Nicole Long.

### Materials availability

This study did not generate new unique reagents.

### Data and code availability

The datasets generated in the current study, along with all experimental codes used for data collection and data analysis will be made publicly available via the Long Term Memory Lab website (http://longtermmemorylab.com/publications/) upon publication.

## Acknowledgments

This work was supported by a grant from the National Institutes of Health (NINDS R01 NS132872, PI: NML).

## Author contributions

Devyn E. Smith: Conceptualization; Data curation; Formal analysis; Project administration; Software; Visualization; Writing—Original draft; Writing—Review & editing. Nicole M. Long: Conceptualization; Formal analysis; Funding acquisition; Project administration; Software; Supervision; Visualization; Writing—Original draft; Writing—Review & editing.

## Star Methods

### Participants

Seventy six native English speakers from the University of Virginia community participated, with thirty eight participants enrolled in each experiment (E1: 28 female; age range = 18-32, mean age = 20.47 years; E2: 26 female; age range = 18-32, mean age = 20.5 years). All participants had normal or corrected-to-normal vision. Informed consent was obtained in accordance with University of Virginia Institutional Review Board for Social and Behavioral Research and participants were compensated for their participation. Our sample size was determined *a priori* based on behavioral pilot data (E1, N = 5; E2, N = 3) described in the pre-registration report of this study (https://osf.io/tfq9u). A total of four participants (two each from E1 and E2) were excluded from the final dataset due to EEG event markers not being recorded. Thus data are reported for the remaining seventy two participants. These data have previously been reported^21^; all of the analyses and results described here are novel.

### Recognition Task Experimental Design

We conducted two recognition memory experiments (E1, E2) and manipulated test-phase instructions between subjects. Participants’ goal was to successfully retrieve study items (E1) or to detect new items (E2). Stimuli consisted of 1602 words, drawn from the Toronto Noun Pool^52^. From this set, 640 words were randomly selected for each participant. Participants completed four phases. Phase 3 was divided into two subsets; the practice subset preceded Phase 1 and the main subset preceded Phase 3.

*Phase 1.* In each of 10 runs, participants viewed a list containing 16 words, yielding a total of 160 trials. On each trial, participants saw a single word presented for 2000 ms followed by a 1000 ms inter-stimulus interval (ISI). In E1, participants were instructed to study the presented word in anticipation of a later memory test and did not make any behavioral responses. In E2, participants were instructed to read the words silently and did not make any behavioral responses.

*Phase 2.* Participants completed a recognition memory test with different memory goals. On each trial, participants viewed a word which had either been presented during Phase 1 (target) or had not been presented (lure). In E1, participants’ task was to make an old or new judgement for each word using a confidence rating scale from 1 to 4, with 1 being definitely new and 4 being definitely old. In E2, the task was framed as a detection phase in which participants’ task was to detect new words that were not presented in Phase 1. Participants made an old or new judgment without the use of a confidence rating scale for each word by pressing one of two buttons (“d” or “k”). Response mappings were counterbalanced across participants. Phase 2 trials were self-paced and separated by a 1000 ms ISI. There were a total of 320 test trials with all 160 Phase 1 words presented along with 160 novel lures.

*Phase 3.* Prior to beginning Phase 1, participants completed three practice runs of a flanker task in which they made speeded responses to a central target in a string of congruent (e.g. *>>>>>>>*) or incongruent (e.g. *<<<><<<*) arrows. Feedback was presented immediately after each response as either a green check mark indicating a correct response or a red X indicating an incorrect response. Response duration, the interval in which a response was accepted, was initially set to 375 ms based on pilot data. To maintain difficulty and ensure an approximately balanced number of correct and incorrect responses during the main subset of Phase 3, response duration was individually adjusted based on participants’ accuracy following each practice run. If accuracy was below 50%, response duration increased by 25 ms, if accuracy was above 50%, response duration decreased by 25 ms. Thus, after completing the three practice runs, the final response duration could be a minimum of 300 ms and a maximum of 450 ms. After completing Phase 2, participants completed the main subset of Phase 3 which consisted of six runs of the flanker task. Throughout the main subset of Phase 3, the response duration was fixed to that obtained from the final practice run.

*Phase 4.* Participants completed a final recognition memory test in which all the words from Phase 1 and 2 were presented along with novel lures. Trials were self-paced and participants made old or new judgements for each word using a confidence rating scale from 1 to 4, with 1 being definitely new and 4 being definitely old. Trials were separated by a 1000 ms ISI. There were a total of 640 test trials with all 320 Phase 2 words presented along with 320 novel lures. As our analyses focus on Phases 2 and 3, we do not consider the final test data further.

### No-feedback flanker task

Forty five native English speakers from the University of Virginia community participated (28 female; age range = 18-50, mean age = 21 years). All participants had normal or corrected-to-normal vision. Informed consent was obtained in accordance with University of Virginia Institutional Review Board for Social and Behavioral Research and participants were compensated for their participation. These data were collected as part of another study which dictated the sample size (described in the pre-registration report of the associated study; osf.io/buegm). A total of two participants were excluded from the final dataset based on the assumption that they inverted the response options (accuracy *<* 10%). Thus data are reported for the remaining forty three participants.

The task design was identical to the feedback flanker task described above, except that in place of feedback in the form of a green check or red X, participants were shown a visual mask.

### EEG Data Acquisition and Preprocessing

EEG recordings were collected using a BrainVision system and an ActiCap equipped with 64 Ag/AgCl active electrodes positioned according to the extended 10-20 system. All electrodes were digitized at a sampling rate of 1000 Hz and were referenced to electrode FCz. Offline, electrodes were later converted to an average reference. Impedances of all electrodes were kept below 50kΩ. Electrodes that demonstrated high impedance or poor contact with the scalp were excluded from the average reference calculations; however, these electrodes were included in all subsequent analysis steps. Bad electrodes were determined by voltage thresholding (see below).

Custom python codes were used to process the EEG data. We applied a high pass filter at 0.1 Hz, followed by a notch filter at 60 Hz and harmonics of 60 Hz to each participant’s raw EEG data. We then performed three preprocessing steps^53^ to identify electrodes with severe artifacts. First, we calculated the mean correlation between each electrode and all other electrodes as electrodes should be moderately correlated with other electrodes due to volume conduction. We z-scored these means across electrodes and rejected electrodes with z-scores less than -3. Second, we calculated the variance for each electrode, as electrodes with very high or low variance across a session are likely dominated by noise or have poor contact with the scalp. We then z-scored variance across electrodes and rejected electrodes with a *|*z*| >* = 3. Finally, we expect many electrical signals to be autocorrelated, but signals generated by the brain versus noise are likely to have different forms of autocorrelation. Therefore, we calculated the Hurst exponent, a measure of long-range autocorrelation, for each electrode and rejected electrodes with a *|*z*| >* = 3. Electrodes marked as bad by this procedure were excluded from the average re-reference. We then calculated the average voltage across all remaining electrodes at each time sample and re-referenced the data by subtracting the average voltage from the filtered EEG data. We used wavelet-enhanced independent component analysis^54^ to remove artifacts from eyeblinks and saccades.

### EEG Data Analysis

For both E1 and E2, we applied the Morlet wavelet transform (wave number 6) to the entire EEG time series across electrodes, for each of 46 logarithmically spaced frequencies (2-100 Hz;^55^). After log-transforming the power, we downsampled the data by taking a moving average across 100 ms time intervals from -1000 to 3000 ms relative to the response and sliding the window every 25 ms, resulting in 157 time intervals (40 non-overlapping). Mean and standard deviation power were calculated across all trials and across time points for each frequency. Power values were then z-transformed by subtracting the mean and dividing by the standard deviation power. We followed the same procedure for the flanker task, with 117 overlapping (30 non-overlapping) time windows from 1000 ms preceding to 2000 ms following the response.

### Region of Interest

We examined theta power across two regions of interest (ROIs), left central (FC1, FC3, FC5, C1, C3, C5, CP1, CP3, CP5) and right central (FC2, FC4, FC6, C2, C4, C6, CP2, CP4, CP6). We specifically focus on the frontocentral region as prior work has demonstrated that theta power in these regions dissociates both hits and correct rejections^56–59^ and positive and negative feedback^14, 15, 60^.

### Univariate Analyses

To test the effect of instructions on theta power, our two conditions of interest were hits (correctly recognized targets) and correct rejections (CRs, correctly rejected lures) in Phase 2 in E1 and E2. We compared theta power across hits and CRs separately for each ROI. For each participant, we calculated z-transformed theta power across both ROIs in each of the two conditions, across 100 ms intervals from 500 ms pre-response to 500 ms post-response, as well as averaged over the 500 ms pre-response interval and the 500 ms post-response interval.

### Pattern Classification Analyses

Pattern classification analyses were performed using penalized (L2) logistic regression implemented via the sklearn module (0.24.2) in Python and custom Python code. We used our pilot data to determine classifier features. We compared performance of two pattern classifiers trained to dissociate positive vs. negative feedback trials in the flanker task. We considered using the 500 ms post-feedback interval; however, during this interval, feedback is visually presented (a green check mark for correct trials and a red x for incorrect trials). Therefore, it is possible that the decoder would leverage properties of the visual stimuli instead of feedback-specific responses. To avoid this potential confound, we trained the classifiers on the 100 ms time interval preceding the response as this interval is less contaminated by visual inputs and is likely to capture internal feedback signals^17, 34, 61^. Within participant classifiers trained on spectral signals averaged across 63 electrodes and 46 frequencies yielded higher mean classification accuracy (M = 58.8%, SD = 5.04%) than within participant classifiers trained on spectral signals averaged across the theta frequency band and 20 central electrodes (M = 56.4%, SD = 4.81%). Therefore, for all classification analyses, classifier features were comprised of spectral power across 63 electrodes and 46 frequencies. Before pattern classification analyses were performed, an additional round of z-scoring was performed across features (electrodes and frequencies) to eliminate trial-level differences in spectral power^62–64^. Thus, mean univariate activity was matched precisely across all conditions and trial types. Classifier performance was assessed in two ways. “Classification accuracy” represented a binary coding of whether the classifier successfully guessed the type of flanker feedback, positive or negative. We used classification accuracy for general assessment of classifier performance (i.e., whether feedback could be decoded). “Classifier evidence” was a continuous value reflecting the logit-transformed probability that the classifier assigned the correct feedback label (positive, negative) for each trial. Classifier evidence was used as a trial-specific, continuous measure of feedback information, which was used to assess the degree of positive feedback evidence present following hit and CRs during Phase 2.

### Cross-Study Feedback Classification

To measure feedback evidence in E1 and E2, we conducted three stages of classification using similar methods as in our prior work^21, 23^. First, we conducted within participant leave-one-run-out cross-validated classification (penalty parameter = 1) on all participants who completed the flanker task (N = 72). For each participant, we generated true and null classification accuracy values using classifiers trained and tested on the 100 ms time interval preceding the flanker response. We sub-sampled to equate the number of correct and incorrect trials and excluded trials with no response. We permuted condition labels (positive feedback, negative feedback) for 1000 iterations to generate a null distribution for each participant. Any participant whose true classification accuracy fell above the 90th percentile of their respective null distribution was selected for further analysis (N = 69). Second, we conducted leave-one-participant-out cross-validated classification (penalty parameter = 0.0001) on the selected participants to validate the feedback classifier. We again sub-sampled to ensure that all participants contributed the same number of correct and incorrect trials, meaning that the classifier would not be biased towards any specific participants. We obtain a mean classification accuracy of 59.42% (SD = 9.520%) which is significantly above chance (*t* _68_ = 8.171, *p <* 0.0001, *d* = 1.401), indicating that the cross-study feedback classifier is able to distinguish positive and negative feedback. Finally, we applied the cross-study feedback classifier to the Phase 2 trials of E1 and E2, specifically in 100 ms intervals from 500 ms pre-response to 500 ms post response. We extracted classifier evidence, the logit-transformed probability that the classifier assigned a given Phase 2 trial a label of positive feedback or negative feedback. This approach provides a trial-level estimate of positive feedback evidence during hits and CRs.

We followed the same procedure to measure correct evidence using the no-feedback flanker data. First, we conducted within participant leave-one-run-out cross-validated classification (penalty parameter = 1) on only those participants who had a minimum of 11 trials per condition (N = 19). This minimum was required to match the amount of data used for classification across the feedback and no-feedback experiments. For each participant, we generated true and null classification accuracy values using classifiers trained and tested on the 100 ms time interval preceding the flanker response. We sub-sampled to equate the number of correct and incorrect trials and excluded trials with no response. We permuted condition labels (correct, incorrect) for 1000 iterations to generate a null distribution for each participant. Any participant whose true classification accuracy fell above the 90th percentile of their respective null distribution was selected for further analysis (N = 15). Second, we conducted leave-one-participant-out cross-validated classification (penalty parameter = 0.0001) on the selected participants to validate the feedback classifier. We again sub-sampled to ensure that all participants contributed the same number of correct and incorrect trials, meaning that the classifier would not be biased towards any specific participants. We obtain a mean classification accuracy of 64.85% (SD = 10.16%) which is significantly above chance (*t* _14_ = 5.484, *p* = 0.0001, *d* = 2.083), indicating that the cross-study no-feedback classifier is able to distinguish correct from incorrect responses. Finally, we applied the cross-study no-feedback classifier to the Phase 2 trials of E1 and E2, specifically in 100 ms intervals from 500 ms pre-response to 500 ms post response. We extracted classifier evidence, the logit-transformed probability that the classifier assigned a given Phase 2 trial a label of correct or incorrect. This approach provides a trial-level estimate of correct evidence during hits and CRs.

### Statistical Analyses

We used mixed effects ANOVAs and *t* -tests to assess the effect of experiment (E1, E2), response (hit, CR) and time interval on frontocentral theta power and feedback evidence.

We used paired-sample *t* -tests to compare classification accuracy across participants to chance decoding accuracy, as determined by permutation procedures. Namely, for each participant, we shuffled the condition labels of interest (e.g., “positive” and “negative” for the feedback classifier) and then calculated classification accuracy. We repeated this procedure 1000 times for each participant and then averaged the 1000 shuffled accuracy values for each participant. These mean values were used as participant-specific empirically derived measures of chance accuracy.

