## Supplemental Results for "Successful retrieval is its own reward"

**Supplemental Material for: Successful retrieval is its own reward**

Author List: Devyn E. Smith[1], Nicole M. Long[1*]

Affiliations:

1: Department of Psychology, University of Virginia 22904

The authors declare no competing interests.

### Hits selectively recruit post-response positive feedback/accuracy signals.

We focused specifically on hits and correct rejections (CRs) in the main manuscript given that both response types are correct, but only hits reflect successful retrieval. However, to the extent that either objective "oldness" or subjective "oldness" induce the feedback and accuracy related responses that we observed, either or both misses and false alarms (FAs) could induce positive feedback and correct evidence increases. It is important to note that some participants contributed a small number of error trials. Specifically, participants committed a miss response to on average 42 trials (min = 5, max = 104) in E1 and 47 trials (min = 11, max = 96) in E2. Participants committed a false alarm on average 30 trials (min = 7, max = 111) in E1 and 28 trials (min = 3, max = 71) in E2.

We assessed positive feedback evidence across all four response types in each experiment (Figure 1, top row). We conducted a 4 $\times$ 2 mixed effects ANOVA with factors of response type (hit, miss, CR, FA) and experiment (E1, E2). Average positive feedback evidence over the post-response interval (0-500ms) was the dependent variable. We find a significant main effect of response ($F_{3,210}$ = 6.579, $p = 0.0003$, $\eta_p{}^2$= 0.0859). We do not find a significant main effect of experiment ($F_{1,70}$ = 0.8487, $p = 0.3601$, $\eta_p{}^2$= 0.0120) or interaction between response and experiment ($F_{3,210}$ = 0.7335, $p = 0.5531$, $\eta_p{}^2$= 0.0104). Post-hoc comparisons revealed significantly greater post-response positive feedback evidence for hits (M = 0.0298, SD = 0.0564) relative to both misses (M =-0.0085, SD = 0.0694, $t_{71}$ = 6.173, $p < 0.0001$, $d$ = 0.7275) and false alarms (M = 0.0052, SD = 0.0824, $t_{71}$ = 3.242, $p = 0.0018$, $d$ = 0.3820).

We assessed correct evidence across all four response types in each experiment (Figure 1, bottom row). We conducted a 4 $\times$ 2 mixed effects ANOVA with factors of response type (hit, miss, CR, FA) and experiment (E1, E2). Average correct evidence over the post-response interval (0-500ms) was the dependent variable. We find a significant main effect of response ($F_{3,210}$ = 11.48, $p < 0.0001$, $\eta_p{}^2$= 0.1409). We do not find a significant main effect of experiment ($F_{1,70}$ = 0.3082, $p = 0.5805$, $\eta_p{}^2$= 0.0044) or interaction between response and experiment ($F_{3,10}$ = 1.640, $p = 0.1813$, $\eta_p{}^2$= 0.0229). Post-hoc comparisons revealed significantly greater post-response correct evidence for hits (M = 0.0325, SD = 0.0563) relative to both misses (M = -0.0143, SD = 0.0628, $t_{71}$ = 8.521, $p < 0.0001$, $d$ = 1.004) and false alarms (M = 0.0097, SD = 0.0733, $t_{71}$ = 3.243, $p = 0.0018$, $d$ = 0.3822).

### Greater pre-response congruency evidence for CRs than hits

Because correct responses (and thus positive feedback) tend to derive from congruent trials whereas incorrect responses (negative feedback) tend to derive from incongruent trials, the positive feedback/correct evidence dissociation that we observe may be due to congruency, and more broadly, response conflict and cognitive control, rather than feedback-related signals. To address this possibility, we developed two congruency training sets, one from each experiment (feedback, no-feedback).

We conducted a multivariate pattern classification analysis in which we trained a classifier to discriminate congruent vs. incongruent trials based on a feature space composed of all 63 electrodes and 46 frequencies ranging from 2-100 Hz. For this analysis, we averaged z-power over the 300 ms following stimulus onset. Using leave-one-participant-out cross-validated classification (penalty parameter = 0.0001), mean classification accuracy was significantly greater than chance for both the feedback (M = 57.89%, SD = 4.320%; $t_{51}$ = 13.04, $p < 0.0001$, $d$ = 2.586; Figure 2A, top row) and no-feedback (M = 58.25%, SD = 3.710%; $t_x$ = 9.945, $p < 0.0001$, $d$ = 3.145; Figure 2A, bottom row) experiment.

We measured congruent evidence across ten 100 ms time intervals from 500 ms preceding to 500 ms following the test-phase response in E1 and E2 (Figure 2B,C, top row). To specifically test for a pre- vs. post-response dissociation in congruent evidence for hits and CRs across E1 and E2, we averaged signals within the 500 ms pre- and post-response time intervals (Figure 2B,C insets, top row). We conducted a $2 \times 2 \times 2$ mixed effects ANOVA with factors of experiment (E1, E2), time interval (pre-response, post-response) and response type (hit, CR). We report the results of this ANOVA in Table 1 and highlight the key findings here. We find a significant interaction between time interval and response type driven by significantly greater congruent evidence for CRs (M = 0.0128, SD = 0.0400) relative to hits (M = -0.0053, SD = 0.0390) in the pre-response time interval ($t_{71}$ = 3.252, $p = 0.0018$, $d$ = 0.4573) and no significant difference in congruent evidence for CRs (M = 0.0016, SD = 0.0468) relative to hits (M = -0.0007, SD = 0.0505) in the post-response time interval ($t_{71}$ = 0.4578, $p = 0.6485$, $d$ = 0.0466). The three-way interaction between response type, time interval, and experiment was not significant. Bayes factor analysis revealed that a model without the three-way interaction term ($H_0$) is preferred to a model with the three-way interaction term ($H_1$; $BF_{10}$ = 0.29, moderate evidence for $H_0$).

We repeated the congruency analysis using training data from the no-feedback experiment (Figure 2B,C insets, bottom row). We conducted the same mixed effects ANOVA as with the feedback experiment data 1. We find a significant interaction between time interval and response type driven by significantly greater congruent evidence for CRs (M = 0.0038, SD = 0.0422) relative to hits (M = -0.0070, SD = 0.0439) in the pre-response time interval ($t_{71}$ = 2.151, $p = 0.0349$, $d$ = 0.2534) and significantly greater congruent evidence for hits (M = 0.0225, SD = 0.0516) relative to CRs (M = 0.0094, SD = 0.0539) in the post-response time interval ($t_{71}$ = 2.160, $p = 0.0341$, $d$ = 0.2546). The three-way interaction between response type, time interval, and experiment was not significant. Bayes factor analysis revealed that a model without the three-way interaction term ($H_0$) is preferred to a model with the three-way interaction term ($H_1$; $BF_{10}$ = 0.28, moderate evidence for $H_0$).

Together, these analyses reveal more congruent evidence for CRs than hits in the pre-response interval. Thus, we do not find evidence to suggest that the observed post-response positive feedback/correct evidence for hits is due to the increased rate of accurate responses on congruent trials.

**Table 1. Mixed effects ANOVAs assessing congruent evidence as a function of experiment, response type, and time interval for each experiment (feedback, no-feedback).**

| | | Feedback | | | No-Feedback | | |
|---|---|---|---|---|---|---|---|
| Effect | df | *F* | *p* | ${\eta_p}^2$ | *F* | *p* | ${\eta_p}^2$ |
| Main effect of experiment | (1,70) | 0.0066 | 0.9353 | 0.0001 | 0.4526 | 0.5033 | 0.0064 |
| Main effect of response type | (1,70) | **5.140** | **0.0265** | **0.0684** | 0.0657 | 0.7984 | 0.0009 |
| Main effect of time interval | (1,70) | 0.2304 | 0.6327 | 0.0033 | 6.487 | 0.0131 | 0.0848 |
| Interaction of experiment × response type | (1,70) | 0.3685 | 0.5458 | 0.0052 | 1.161 | 0.2849 | 0.0163 |
| Interaction of experiment × time interval | (1,70) | 2.744 | 0.1021 | 0.0377 | 3.079 | 0.0837 | 0.0421 |
| Interaction of response type × time interval | (1,70) | **7.952** | **0.0062** | **0.1020** | **13.83** | **0.0004** | **0.1649** |
| Interaction of experiment × response type × time interval | (1,70) | 0.5677 | 0.4537 | 0.0080 | 0.4141 | 0.5220 | 0.0059 |

*Note*: bold values indicate $p < 0.05$.

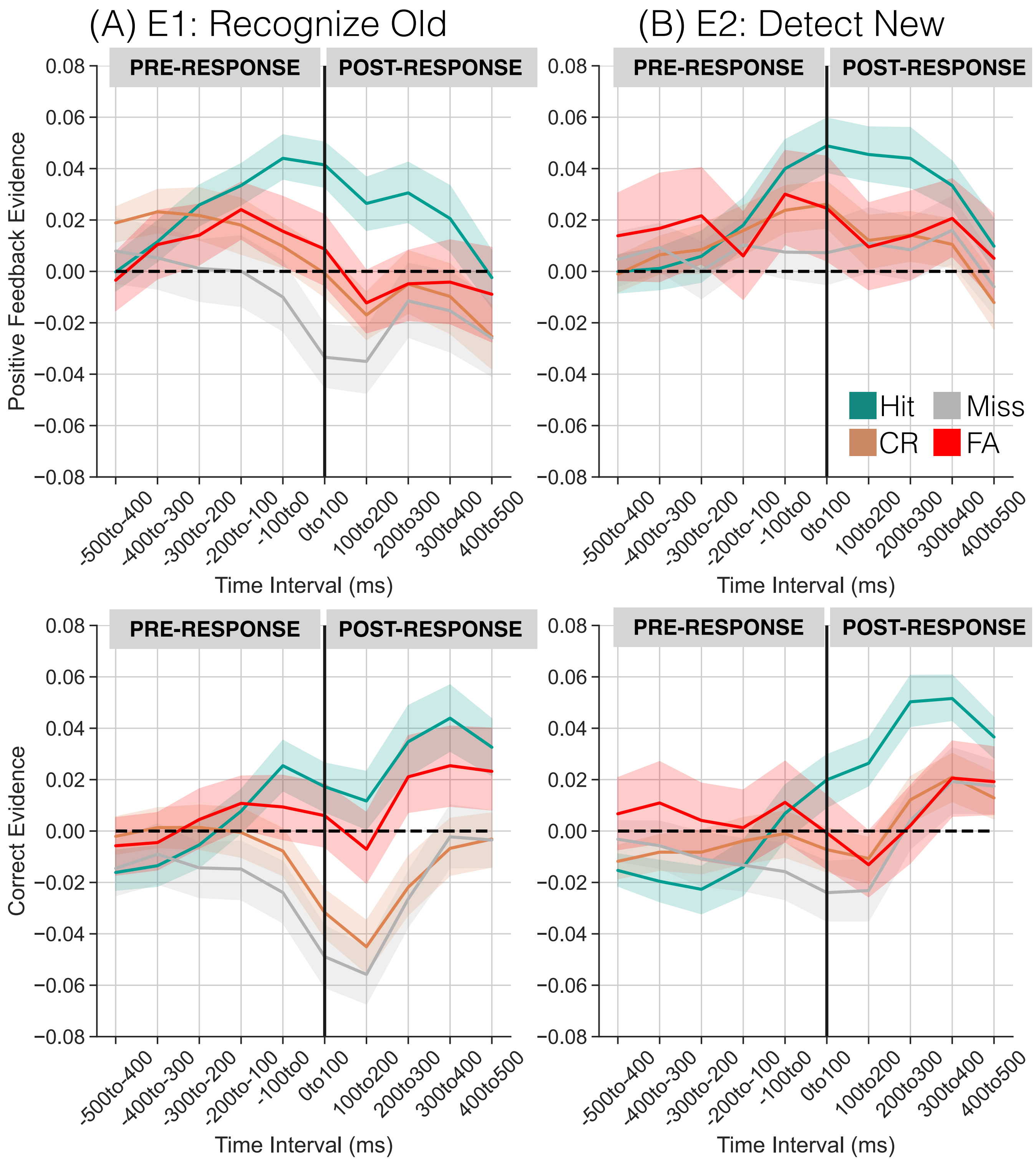


Figure 1. **Positive feedback and correct evidence preceding and following memory responses in E1 and E2.** The top row shows results from the feedback trained classifier; the bottom row shows results from the no-feedback trained classifier. Positive y-axis values indicate greater positive feedback/correct evidence. The solid vertical line at time 0 to 100 ms indicates the response. Each panel shows positive feedback/correct evidence separated by response type (teal: hit; orange: CR; grey, Miss; red; False alarm) across the 500 ms pre-response and post-response time intervals. There is greater post-response positive feedback/correct evidence for hits compared to both misses and false alarms ($p$'s $< 0.01$), regardless of experiment goals. Error bars represent standard error of the mean.

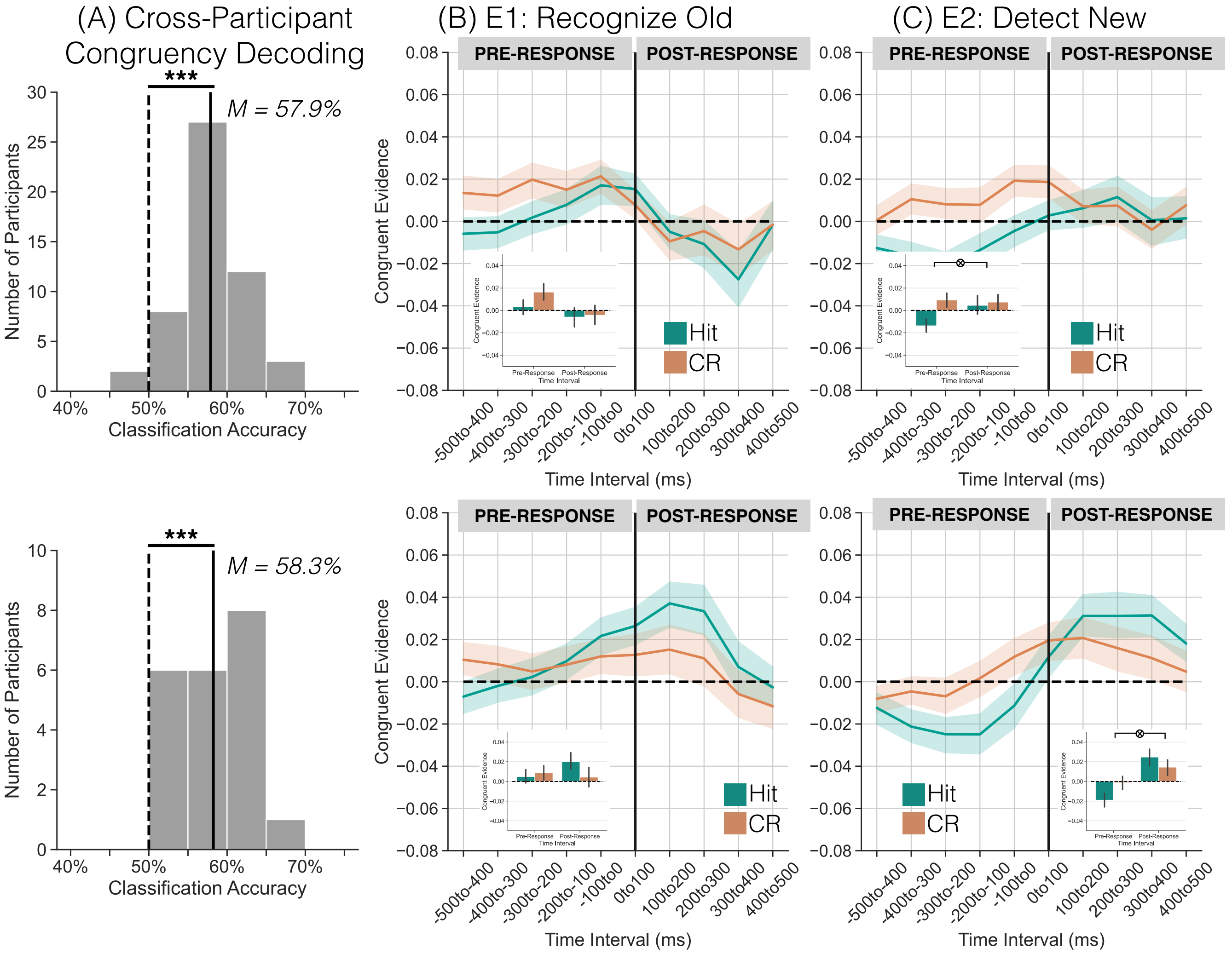


Figure 2. **Congruent evidence preceding and following memory responses in E1 and E2. (A)** Mean classification accuracy across all participants (solid vertical line) is shown along with a histogram of classification accuracies for individual participants (gray bars) and mean classification accuracy for permuted data across all participants (dashed vertical line). Mean classification accuracy was 57.9% for the feedback experiment (top row) and 58.3% for the no-feedback experiment (bottom row), both of which differed significantly from chance (two-tailed, paired t-tests, $p < 0.0001$). **(B-C)** Positive y-axis values indicate greater congruent evidence. The solid vertical line at time 0-100 ms indicates the response. Each panel shows congruent evidence separated by response type (teal: hit; orange: CR) across the 500 ms pre-response and post-response time intervals. There is greater pre-response congruent evidence for CRs compared to hits across both experiments ($p$'s $< 0.05$). Error bars represent standard error of the mean.
